# Is Palmer’s elm leaf goldenrod real? The Angiosperms353 kit provides within-species signal in *Solidago ulmifolia* s.l

**DOI:** 10.1101/2021.01.07.425781

**Authors:** James B. Beck, Morgan L. Markley, Mackenzie G. Zielke, Justin R. Thomas, Haley J. Hale, Lindsay D. Williams, Matthew G. Johnson

**Affiliations:** Department of Biological Sciences, Wichita State University, Wichita, KS 67260 U.S.A; Botanical Research Institute of Texas, Fort Worth, Texas 76107 U.S.A; NatureCITE, Springfield, Missouri 65803 U.S.A; Department of Biological Sciences, Texas Tech University, Lubbock, Texas 79409 U.S.A

**Keywords:** Angiosperms353, hyb-seq, *Solidago*, *Solidago ulmifolia* var. *palmeri*, species delimitation

## Abstract

The genus *Solidago* represents a taxonomically challenging group due to its sheer number of species, putative hybridization, polyploidy, and shallow genetic divergence among species. Here we use a dataset obtained exclusively from herbarium specimens to evaluate the status of *Solidago ulmifolia* var. *palmeri,* a morphologically subtle taxon potentially confined to Alabama, Arkansas, Mississippi, and Missouri. A multivariate analysis of both discrete and continuous morphological data revealed no clear distinction between *S. ulmifolia* var. *palmeri* and *Solidago ulmifolia* var. *ulmifolia. Solidago ulmifolia* var. *palmeri*’s status was also assessed with a phylogenomic and SNP clustering analysis of data generated with the “Angiosperms353” probe kit. Neither analysis supported *Solidago ulmifolia* var. *palmeri* as a distinct taxon, and we suggest that this name should be discarded. The status of *Solidago delicatula* (formerly known as *Solidago ulmifolia* var. *microphylla)* was also assessed. Both morphological and phylogenic analyses supported the species status of *S. delicatula* and we suggest maintaining this species at its current rank. These results highlight the utility of the Angiosperms353 probe kit, both with herbarium tissue and at lower taxonomic levels. Indeed, this is the first study to utilize this kit to identify genetic groups within a species.

---

Many botanists find *Solidago* L. taxonomically challenging (Fernald 1950; Croat 1967; Correll and Johnston 1970; Voss 1996; Nesom 1993; Zhang 1996; Cook 2002), a problem stemming from the sheer number of species involved, putative hybridization, and polyploidy. Unfortunately, first-generation DNA sequence data have been of little use in clarifying *Solidago* species boundaries due to the low observed genetic diversity in the genus. Most notable are three DNA barcoding studies. Among the eight groups examined in Kress et al. (2005), *Solidago* harbored the lowest level of diversity at 10 highly variable sequence loci, exhibiting no substitutions at the “universal” barcoding region *psbA-trnH.* Fazekas et al. (2008, 2009) then examined nine potential barcoding regions in 32 genera and commented that *Solidago* was one of the two most “intractable” genera. Any species delimitation or phylogeny reconstruction in *Solidago* will therefore require genomic datasets, which have shown promise in the genus (Beck and Semple 2015, Jordon-Thaden et al. 2020).

Among the many taxonomic issues in *Solidago* L. is the status of Palmer’s elm leaf goldenrod *(Solidago ulmifolia* Muhlenberg ex Willdenow var. *palmeri* Cronquist). This taxon is distinguished by densely-pubescent stems below the inflorescence (Cronquist 1947), as *Solidago ulmifolia* Muhlenberg ex Willdenow var. *ulmifolia* is viewed as typically glabrous below the inflorescence. As currently circumscribed *S. ulmifolia* var. *palmeri* is relatively common in the Ozark and Boston Mountains of Arkansas/Missouri, with disjunct populations in Mississippi and Alabama (Semple and Cook 2006). Although seemingly distinctive, most regional floras note the presence of some hairs in *S. ulmifolia* var. *ulmifolia-* “glabrous or nearly so” (McGregor et al. 1986, Smith 1994). Additionally, in his description of the taxon Cronquist noted that Alabama *S. ulmifolia* var. *palmeri* material was glabrous in the lower portion of the stem “suggesting a transition to var. *ulmifolia”* (Cronquist 1947). Palmer’s elm leaf goldenrod is part of *Solidago* subsection *Venosae* (G.Don) G.L. Nesom, a group characterized by chiefly cauline/reticulate-veined leaves and secund capitula (Semple and Cook 2006). In this study we combine morphological and genomic datasets to investigate the distinctiveness of Palmer’s elm leaf goldenrod in the context of the larger phylogeny of subsection *Venosae.*

## Materials and methods

### Sampling

All analyses were performed on data obtained exclusively from herbarium specimens-see Appendix 1 for sample information. All samples were either from taxa in which only diploids are known, or were otherwise established as diploid with a chromosome count. Our morphological analyses comprised 114 individuals of *S. ulmifolia* s.l., including 11 *S. delicatula* (formerly known as *Solidago ulmifolia* Muhlenberg ex Willdenow var. *microphylla* A.Gray), 24 *S. ulmifolia* var. *palmeri,* 60 *S. ulmifolia* var. *ulmifolia,* and 19 *Solidago ulmifolia* aff. var. *palmeri* individuals. Given the continuous distribution of stem pubescence we observed, these “aff. var. *palmeri”* individuals were defined as those that exhibited 10-20 hairs along a 3 mm length of the mid-stem. Those with fewer hairs were considered *S. ulmifolia* var. *ulmifolia* and those with more hairs *S. ulmifolia* var. *palmeri*.

For our phylogenetic analyses we included 72 individuals of subsection *Venosae* (Semple and Cook 2006) and outgroups. This sample set included 39 *S. ulmifolia* s.l. individuals [*S. delicatula*, *S. ulmifolia* var. *palmeri*, *S. ulmifolia* var. *ulmifolia*, and *S. ulmifolia* aff. var. *palmeri*]. Sixteen *Solidago rugosa* s.l. individuals were included-note that we didn’t attempt to distinguish among the various taxa in *S. rugosa* s.l. [*S. rugosa* Miller var. *rugosa, S. rugosa* Miller var. *aspera* Fernald*, S. rugosa* Miller var. *celtidifolia* (Small) Fernald, and *Solidago aestivalis* E.P.Bicknell-formerly known as S. *rugosa* Miller var. *sphagnophila* C.Graves.] Four individuals of *Solidago fistulosa* Miller were included. Remaining sampling included taxa placed in an expanded concept of subsection *Venosae* based on a phylogenomic analysis of *Solidago* using 893 nuclear genes (Beck, unpubl. data). This sampling included three specimens of *S. drummondii* Torrey and A.Gray and two taxa formerly placed in *Solidago* subsection *Argutae* (Mackenzie) G.L.Nesom: three individuals of *Solidago brachyphylla* Chapman ex Torrey & A.Gray, and two individuals of *Solidago auriculata* Shuttleworth ex S.F.Blake. *Solidago odora* Aiton was not included as preliminary phylogenomic data indicates that it is distantly related to subsection *Venosae* taxa. Two individuals of *Solidago patula* Muhlenberg ex Willdenow and three individuals of *Solidago uliginosa* Nuttall were included as outgroups.

### Morphological analyses

All morphological analyses of *S. ulmifolia* s.l. presented here were conducted in R version 4.0.0 (R Core Team 2020). For each measured specimen we initially assessed 41 morphological characters (supplementary appendix S1). We first assessed plots of single characters and biplots of two characters with ggplot2 (Wickham 2016) to discover characters that clearly distinguished taxa. A combination of two characters clearly separated *Solidago delicatula*, and further analyses excluded this taxon. After removing two characters which had missing data and all discrete (i.e. “count”) characters, 27 continuous characters remained. Four continuous characters were removed to eliminate highly correlated pairs of characters (Pearson correlations >0.8), retaining characters that loaded more highly on a preliminary Principal Components Analysis (PCA) conducted with the adegenet package (Jombart, 2008). A PCA was then conducted on the remaining 23 continuous characters. A similar workflow was performed on 11 discrete characters (no strong correlations were detected).

### Phylogenomic analyses

DNA extractions followed a standard CTAB protocol modified for 96 well plates (Beck et al. 2012), and a Qubit fluorometer (Life Technologies, Eugene, Oregon) was used to establish DNA concentration for all extracts. Library preparations were performed using the NEBNext Ultra II DNA Library Prep Kit for Illumina with the NEBNext Multiplex Oligos for Illumina (Dual Index Primers Set 1) (NEB, Ipswich, Massachusetts). Library preparation followed the protocol outlined in Saeidi et al. (2018), with 200 ng of input DNA. Note that 41 samples with low library concentrations were re-amplified (Saeidi et al. 2018) with universal Illumina primers prior to the hybridization reaction. Hybridization was performed with the “Angiosperms353” probe kit (Johnson et al. 2019) using the methods outlined in the Hybridization Capture for Targeted NGS protocol (Arbor Biosciences, Ann Arbor, Michigan). Three reactions were conducted initially, with reaction membership designed to correct for varying final Illumina library concentrations. The first pool comprised 9 samples with library qubits < 6.5 ng/μl, adding 10 μl of each library. The second pool comprised 24 samples with library qubits 6.5 – 14.0 ng/μl, adding 8 μl of each library. The third pool comprised 18 samples with library qubits > 14.0 ng/μl, adding 5 μl of each library. All three reactions were pooled in equal-molar ratios and sequenced (paired end 300 bp) on one lane of Illumina MiSeq version 3 chemistry (Illumina, San Diego, California). The final reaction comprising 21 libraries with DNA concentrations 11.4 – 20.0 ng/μl was performed and sequenced as above.

All analyses performed below were conducted on the Beocat High Performance Computing cluster at Kansas State University (Manhattan, Kansas). Following de-multiplexing, adapters and low-quality sequence were removed with Trimmomatic (Bolger et al. 2014). The bioinformatic workflow HybPiper (Johnson et al. 2016) was then used to align reads and establish sample sequences at each gene using representative target sequences from github.com/mossmatters/Angiosperms353. Mafft (Katoh et al. 2002) was used to align sample sequences at each locus, and trimAl (Capella-Gutiérrez 2009) was used to remove sites missing in > 50% of mafft alignments. RAxML (Stamatakis 2014) “best” ML trees and bootstrap values were obtained with the GTRCAT model of sequence evolution. After all nodes with >30% bootstrap support were collapsed, an Astral (Mirarab 2014) species tree was obtained from these collapsed trees. The “intronerate” script (Johnson et al. 2016) was then run to generate “supercontigs” of both intron and exon sequence at each locus, with alignment and tree-building workflows performed as above.

### Genomic SNP analyses

SNP analyses were performed on 36 *S. ulmifolia* s.l. individuals to further search for geographic/taxonomic patterns. One sample (IL67R) was used as a reference for all other samples to map to. HybSeq SNP Extraction Pipeline (scripts available at: https://github.com/lindsawi/HybSeq-SNP-Extraction) was used to process samples. For each sample, read one and read two were mapped to IL67R according the GATK Variant Discovery Best Practices Workflow. Duplicate reads were removed and variant sites were called using GATK in GVCF mode (HaplotypeCaller). GVCF files were combined and GATK joint genotype caller was used to identify and filter SNPs in sample sequences. SNPs were removed if it were determined they fell below a hard quality filter “QD < 5.0 || FS > 60.0 || MQ < 40.0 || MQRankSum < −12.5 || ReadPosRankSum < −8.0”. Using PLINK, variants were additionally filtered to remove SNPs containing missing data and to reduce the dataset to exclude SNPs with evidence of linkage using PLINK filter “--indep 50 5 2” as a sliding window to assess linkage. Using the unlinked SNP data, eigenvectors were generated for 20 coordinate PCA axes with PLINK. The unlinked SNP file was additionally used to generate ancestry information about the sample population using the package LEA (Frichot and François, 2015). LEA was used to determine the ancestry coefficients of the sample population using lowest cross-entropy. Values for K (number of clusters) was set between 1 and 10, with 50 iterations at each value of K. The appropriate value of K was determined using the slope of the differences between adjacent K-values. The Q-Matrix was used to visualize admixture frequencies for each sample. Pie charts that are representative of the admixture within each sample were then plotted using geographic coordinates.

## Results

### Morphological Analyses

The combination of the number of hairs along the middle leaf abaxial surface midvein (<3 hairs) and upper stem pubescence (<3 hairs) clearly separated *Solidago delicatula* from other members of *S. ulmifolia* s.l. (Fig. 1). In the analysis of 23 continuous characters from the remaining *S. ulmifolia* s.l. taxa, Principal Component 1 (PC1) explained 15.0% of the variation, while Principal Component 2 (PC2) explained 14.0% of the variation. In the analysis of 11 discrete characters from this dataset, PC1 explained 30.2% of the variation, while PC2 explained 12.9% of the variation. Analysis of both discrete and continuous variables failed to recover strong distinctions among *S. ulmifolia* var. *palmeri, S. ulmifolia* var. *ulmifolia,* and *S. ulmifolia* aff. var. *palmeri.* (Fig 2).

**Fig. 1.**
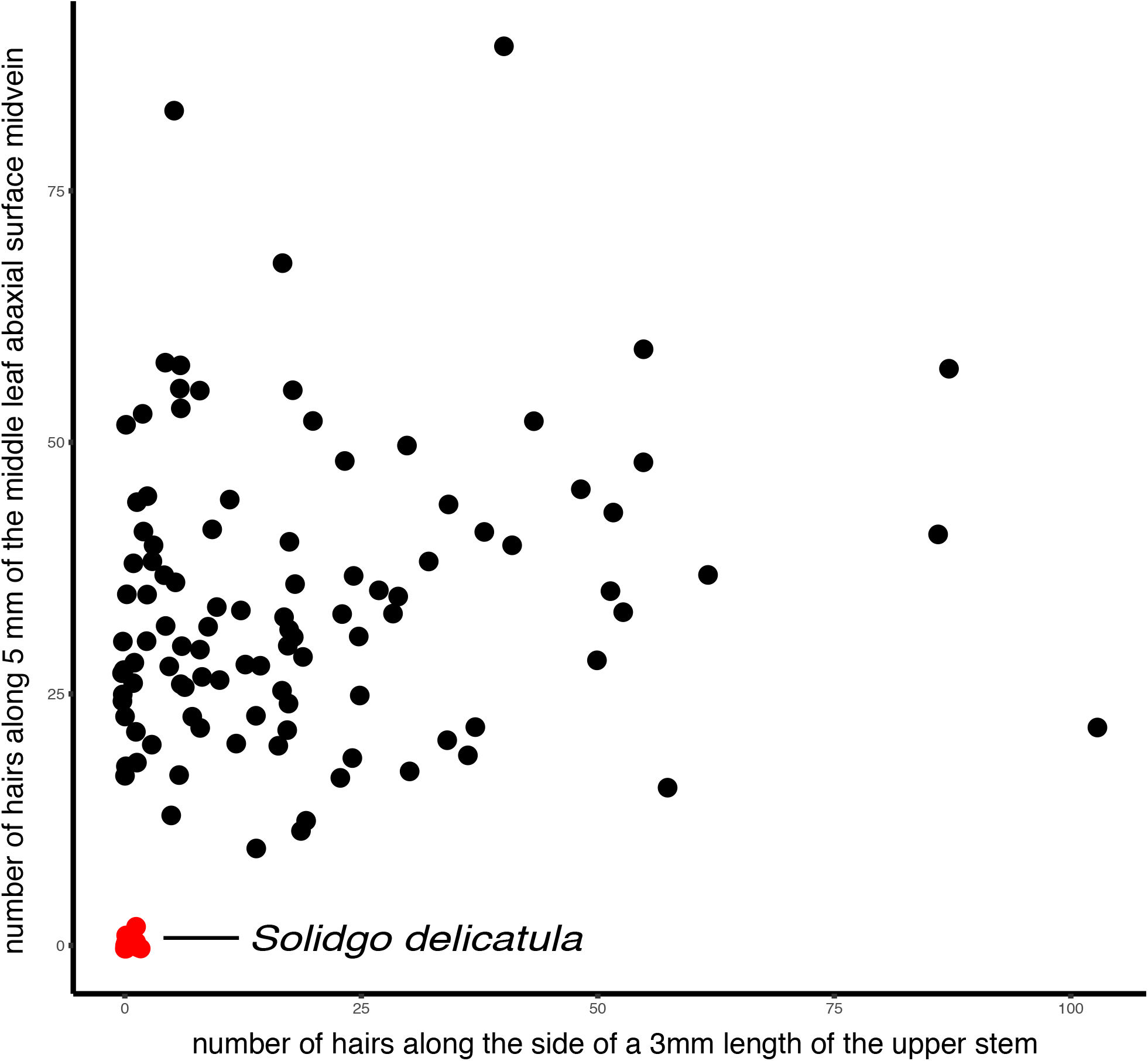
Biplot of two characters (the number of hairs along the middle leaf abaxial surface midvein and upper stem pubescence) that conclusively diagnose *Solidago delicatula* relative to other members of *Solidago ulmifolia* s.l.

**Fig. 2.**
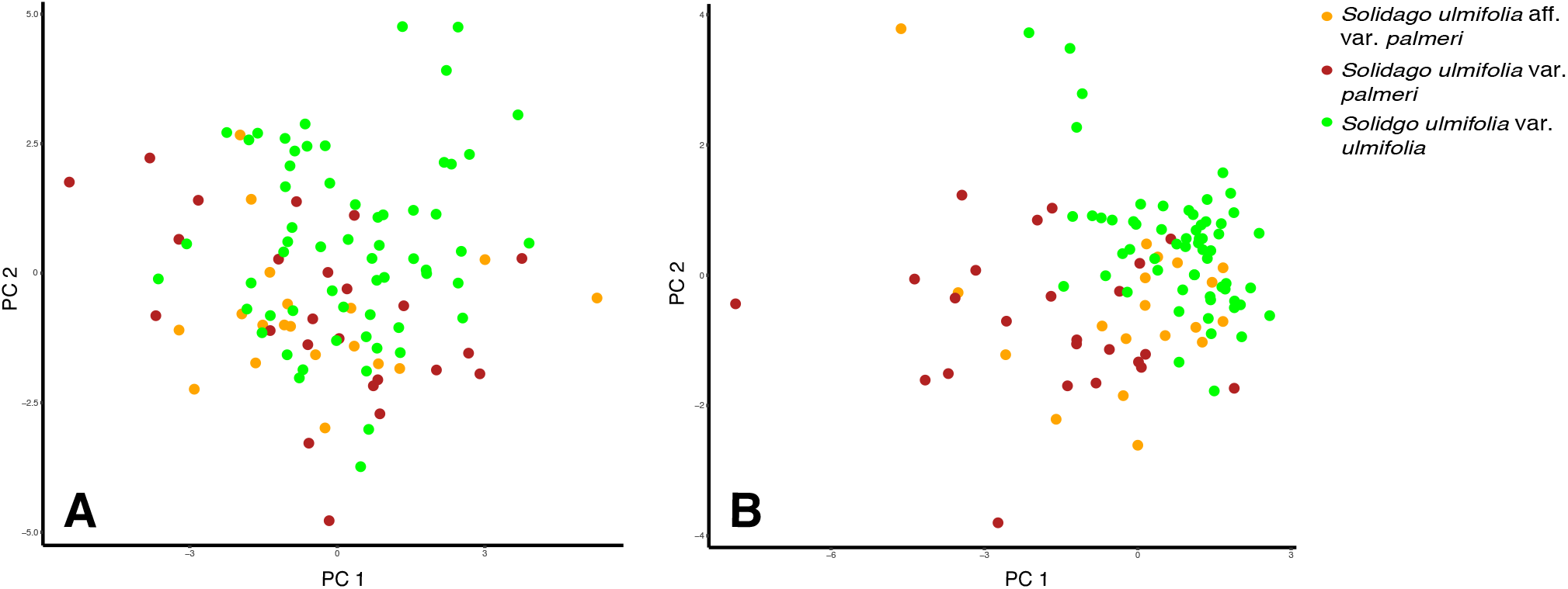
Principal components analysis of morphological variation among taxa. A. Plot of the first two principal components resulting from analysis of 23 continuous characters in *Solidago ulmifolia* var. *palmeri*/*Solidago ulmifolia* aff. var. *palmeri*/*Solidago ulmifolia* var. *ulmifolia* individuals. B. Plot of the first two principal components resulting from analysis of 11 discrete characters in *S. ulmifolia* var. *palmeri*/ *S. ulmifolia* aff. var. *palmeri*/*S. ulmifolia* var. *ulmifolia* individuals.

### Phylogenomic analyses

Data was recovered from 69/72 samples and at 344/353 genes in the intronerate “supercontig” analysis. An average of 143.6 bp of sequence was recovered per gene, and an average of 49,398 bp per sample. There was a significant negative relationship between sequencing success (average gene length) and specimen age *(R*^2^ = 0.197; *P* < 0.001). Astral analysis of 344 nuclear genes identified clades corresponding to all sampled species, with the exception of a single sample of *S. ulmifolia* aff. var. *palmeri* which was not placed in the *S. ulmifolia* s.l. clade (Fig. 3). The *Solidago rugosa* s.l. clade was placed as sister to *S. fistulosa* (99%), with the *S. ulmifolia* s.l. clade sister to a *S. auriculata/S. brachyphylla* clade (64%). These sister relationships were also seen in the genus-wide phylogenomic analysis of *Solidago* using 893 nuclear genes (Beck, unpubl. data), except that in that analysis, *S. ulmifolia* s.l. was sister to *S. brachyphylla,* a clade which was in turn sister to *S. auriculata.* Within the *S. ulmifolia* s.l. clade, although six of the seven *S. delicatula* samples formed a clade (62%), no clades were observed corresponding to *S. ulmifolia* var. *palmeri, S. ulmifolia* aff. var. *palmeri,* or *S. ulmifolia* var. *ulmifolia* (Fig. 3).

**Fig. 3.**
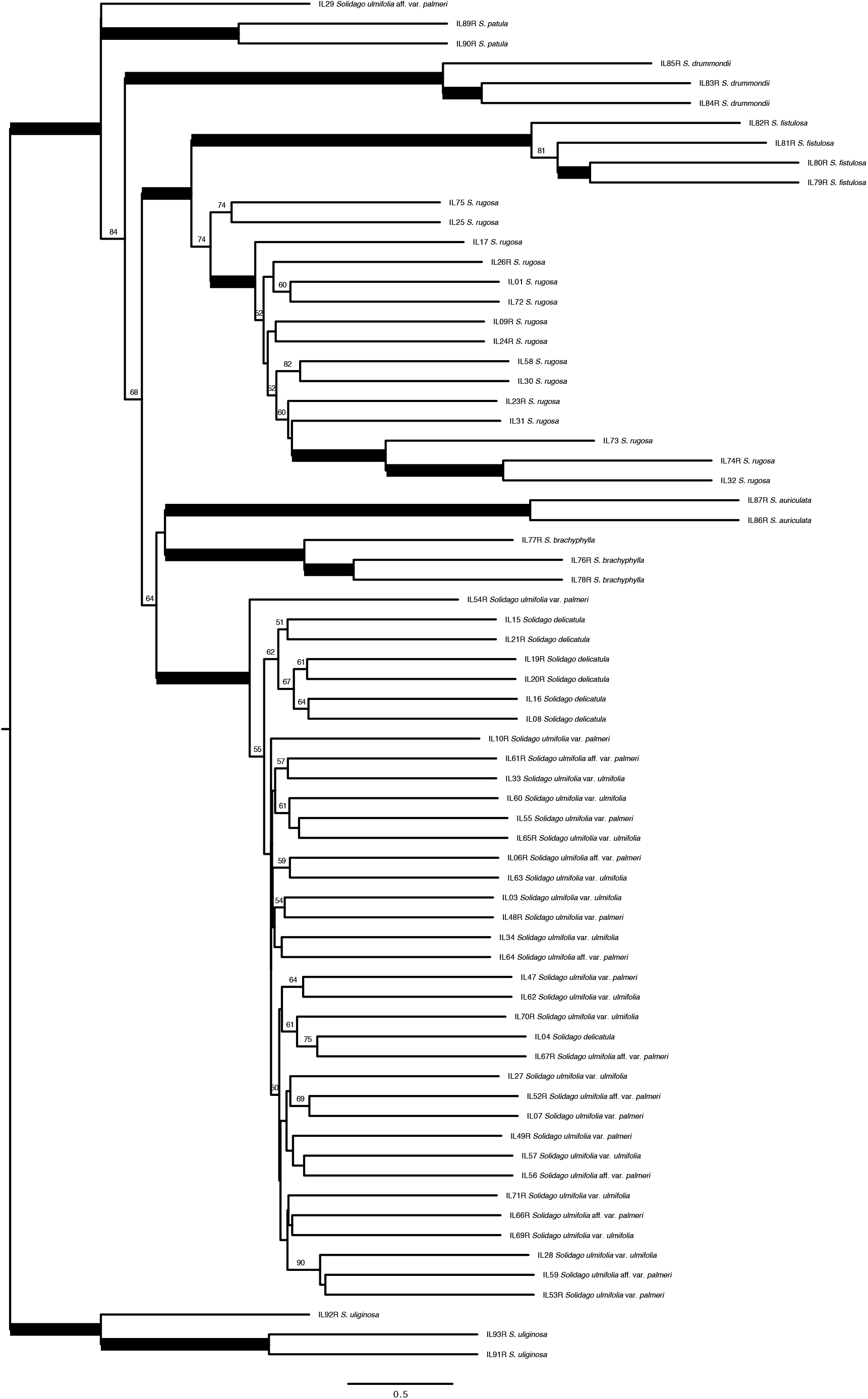
Astral consensus tree resulting from analysis of 344 genes from 69 *Solidago* subsection *Venosae* individuals and outgroups. Nodes with >95% Astral bootstrap support are in bold, otherwise bootstrap values >50% are shown. IL numbers refer to Illumina library numbers which serve as unique identifiers for each sample-see Appendix 1.

### Genomic SNP analyses

A total of 46363 SNPs passed basic filtering based on base quality and coverage depth. Of these, 10595 SNPs had no missing data and of these 4183 passed a check for linkage disequilibrium and were considered unlinked. Using this pruned and unlinked dataset, the lowest clustering entropy was observed with K=3, and ancestry coefficients at this K-value are shown for all samples (Fig. 4). A plot of individuals at the first two principal components, with individuals colored by the maximum LEA Q-matrix cluster score, shows clear separation of these three groups (Fig. 5). A plot of the geographic distribution of each sample colored by their ancestry coefficients (Fig. 6) shows clear geographic structure, with one cluster comprising *S. ulmifolia* var. *palmeri*, *S. ulmifolia* aff. var. *palmeri*, and *S. ulmifolia* var. *ulmifolia* samples found primarily east of the Mississippi River (Fig. 5 cluster 1), a second cluster of *S. ulmifolia* var. *palmeri*, *S. ulmifolia* aff. var. *palmeri*, and *S. ulmifolia* var. *ulmifolia* found primarily in AR/MO (Fig. 5 cluster 2), and a third cluster found primarily in *S. delicatula* samples from AR/KS/OK (Fig. 5 cluster 3).

**Fig. 4.**
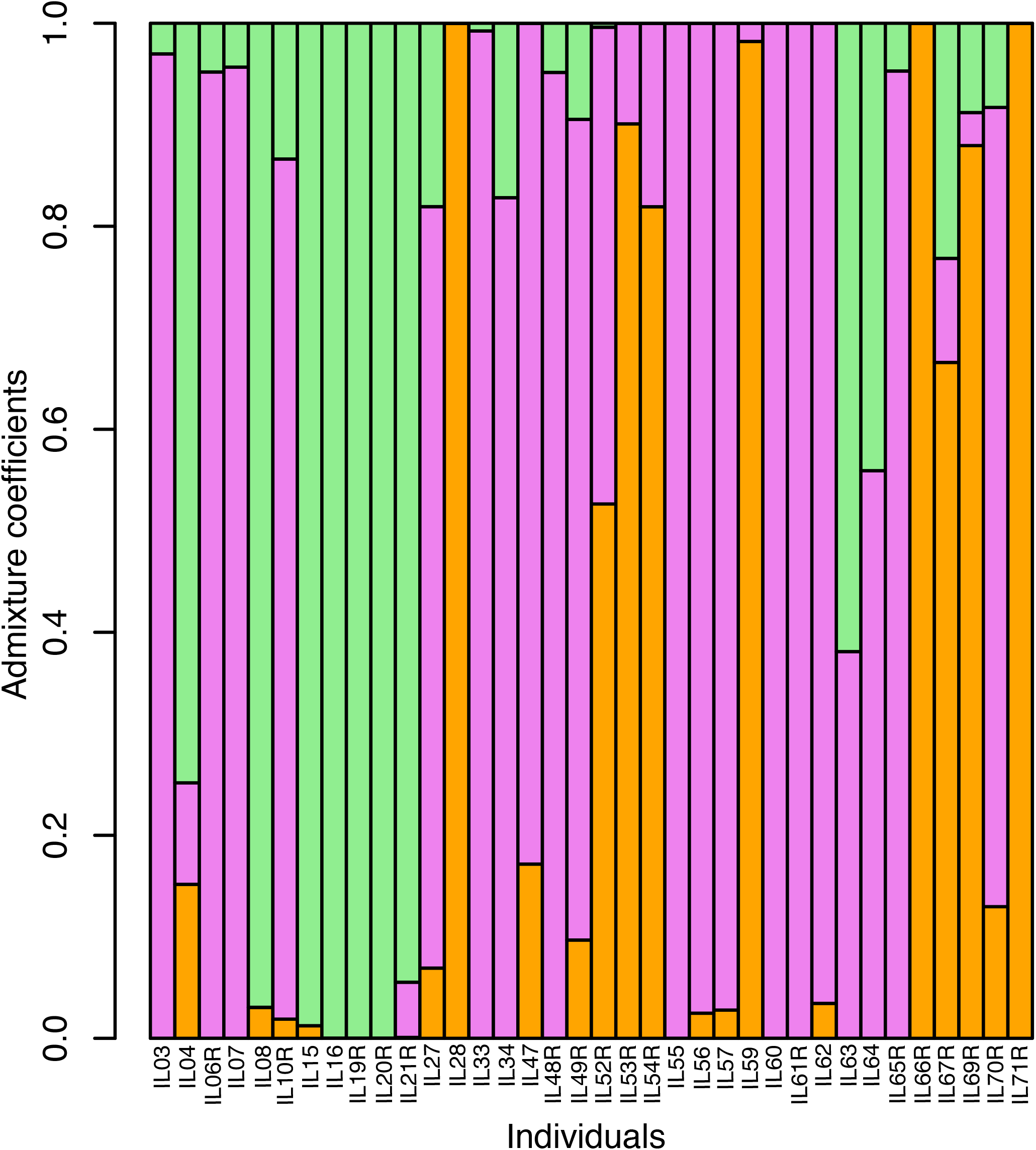
Plot of ancestry coefficients resulting from a LEA analysis of genomic SNPs from 36 *Solidago ulmifolia* s.l. individuals (K=3). IL numbers refer to Illumina library numbers which serve as unique identifiers for each sample-see Appendix 1.

**Fig. 5.**
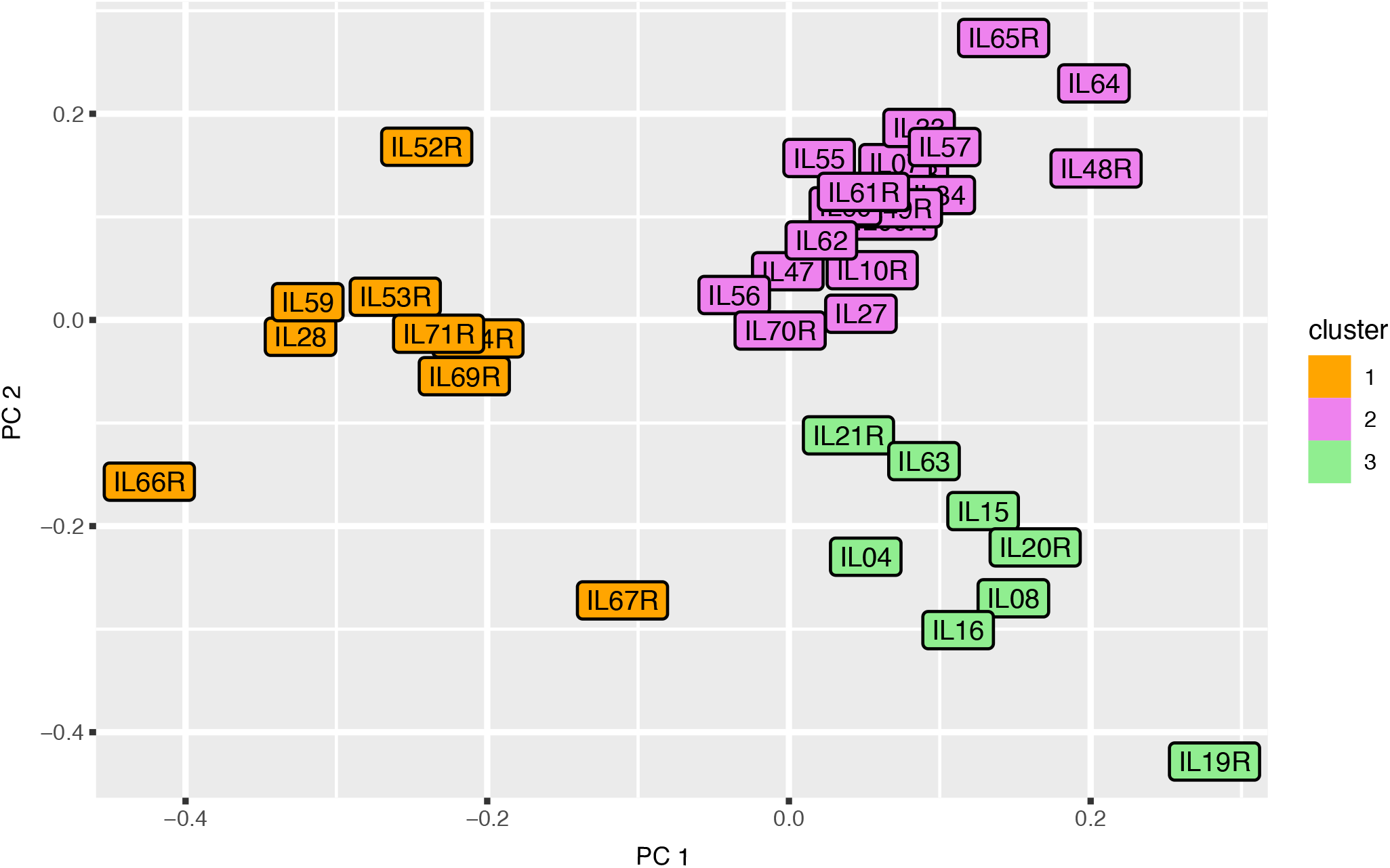
Principal components analysis of SNPs from *Solidago ulmifolia* s.l. individuals, colored according to each sample’s maximum LEA ancestry coefficient. IL numbers refer to Illumina library numbers which serve as unique identifiers for each sample-see Appendix 1.

**Fig. 6.**
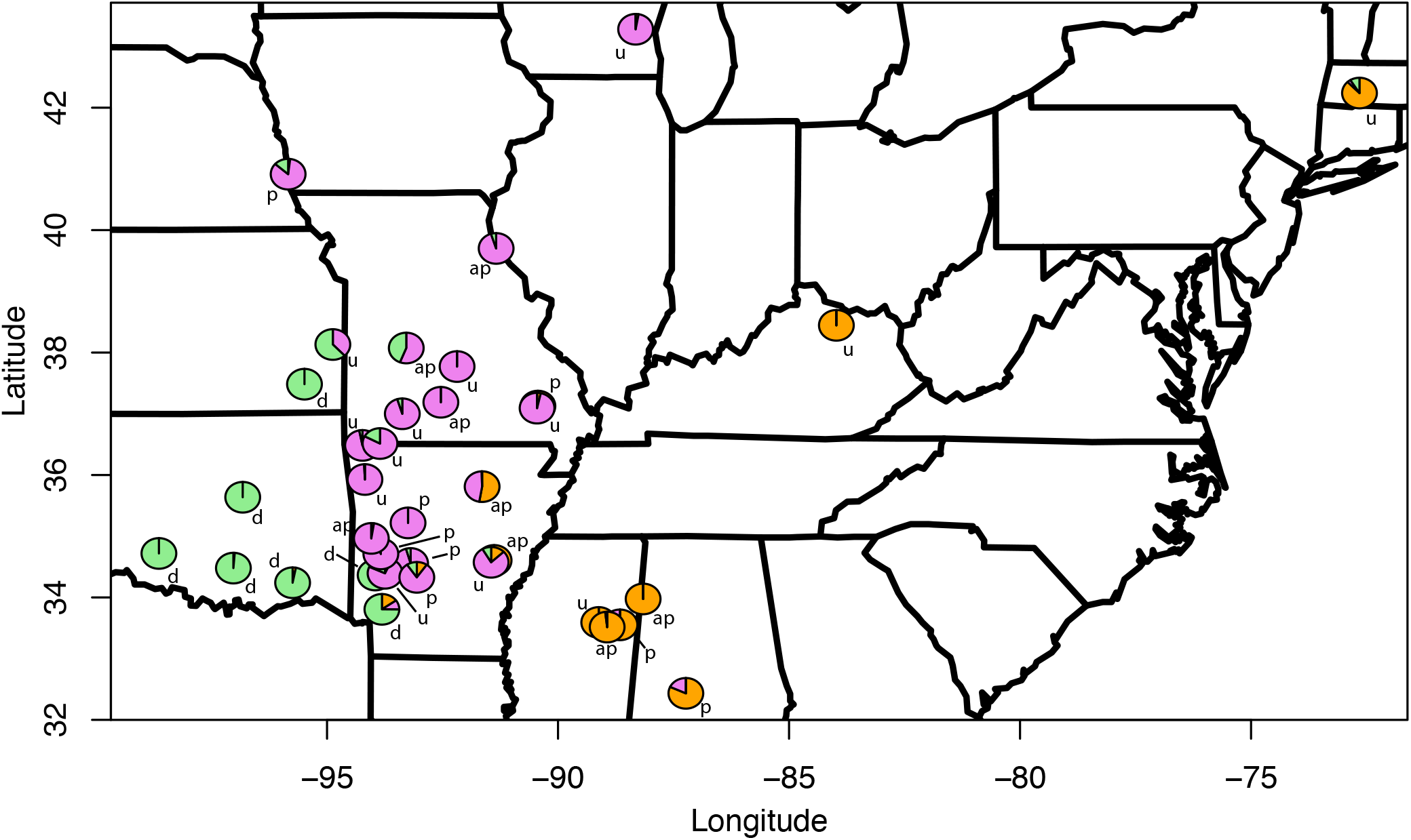
Geographic distribution of 36 *Solidago ulmifolia* s.l. individuals included in the genomic SNP analysis, with pie charts indicating ancestry coefficients from the LEA analysis. Letters denote sample identification (p = *S. ulmifolia* var. *palmeri*, u = *S. ulmifolia* var. *ulmifolia*, ap = *S. ulmifolia* aff. var. *palmeri*).

## Discussion

### Species status

Both morphological and molecular data supported the species status of *Solidago delicatula* (formerly known as *Solidago ulmifolia* Muhlenberg ex Willdenow var. *microphylla).* Two characters (the number of hairs along the middle leaf abaxial surface midvein and upper stem pubescence) clearly separated the 11 *Solidago delicatula* individuals from the remaining *S. ulmifolia* s.l. individuals in a biplot (Fig. 1). Furthermore, 6/7 sequenced *Solidago delicatula* individuals formed a moderately supported (62%) clade (Fig. 3) and all seven *Solidago delicatula* individuals were part of the 8-sample “cluster three” (Figs. 5, 6). *Solidago delicatula* therefore exhibits both morphological and phylogenetic distinctiveness, two species criteria under the General Lineage Concept of Species (de Queiroz 1998). This is in line with contemporary treatments of *Solidago,* which typically recognize *S. delicatula* (Semple and Cook 2006). On the contrary, neither morphological nor molecular data supported the taxonomic status of *S. ulmifolia* var. *palmeri.* Analysis of continuous (Fig. 2A) and discrete (Fig. 2B) morphological data identified at best a morphological cline between *S. ulmifolia* var. *palmeri* and *S. ulmifolia* var. *ulmifolia*, regardless of how *S. ulmifolia* aff. var. *palmeri* individuals are considered. Although *S. ulmifolia* s.l. was identified as almost completely monophyletic in the phylogenomic analysis, no supported clades were identified that corresponded to either *S. ulmifolia* var. *palmeri* or *S. ulmifolia* var. *ulmifolia* (Fig. 3). The genomic SNP analysis provided further clarification. Rather than morphology, clusters one and two corresponded largely to geography, with one cluster comprising *S. ulmifolia* var. *palmeri*, *S. ulmifolia* aff. var. *palmeri*, and *S. ulmifolia* var. *ulmifolia* samples found primarily east of the Mississippi River (Figs. 5, 6), with cluster two comprising *S. ulmifolia* var. *palmeri*, *S. ulmifolia* aff. var. *palmeri*, and *S. ulmifolia* var. *ulmifolia* found primarily in MO/AR (Figs. 5, 6). The *S. ulmifolia* var. *palmeri* morphotype therefore appears to simply represent the presence of genetic variation for increased stem pubescence in certain portions of the *S. ulmifolia* range, and our morphological concept of *S. ulmifolia* should be expanded to accommodate this variation. We therefore suggest discarding the widely used name *S. ulmifolia* var. *palmeri* both in treatments and in herbarium organization. This will not affect the protected status of this taxon, as it is not listed in any of the states in which it occurs.

### Utility of the Angiosperms353 probe kit

Beyond their taxonomic implications, our results further highlight the utility of the Angiosperms353 probe kit, both with herbarium tissue and at lower taxonomic levels. Although sequencing success did drop with increasing specimen age, we were able to recover large datasets from 69/72 samples, with collection years ranging from 1892-2011 (Appendix 1). This joins the growing number of studies that have successfully used this probe kit with herbarium tissue (Brewer et al. 2019; Shee et al. 2020). Considerable signal was also evident at lower taxonomic levels. The monophyly of *Solidago* species were strongly supported (Fig. 3). If one putative interspecific hybrid (IL29) is put aside, all eight included species were recovered as monophyletic, 7/8 with maximum Astral bootstrap support. This is notable given the low level of sequence divergence reported among *Solidago* species in earlier barcoding studies (Kress et al. 2005; Fazekas et al. 2008, 2009). Furthermore, the SNP analysis identified genomic clusters that largely corresponded to both taxonomic and geographic groups within *S. ulmifolia* s.l. (Figs. 5, 6). Although this study joins a growing number that have successfully used this probe kit among shallowly diverged taxa (Larridon et al. 2020; Shee et al. 2020), to our knowledge is the first study to utilize the Angiosperms353 probe kit to identify genetic groups *within* species. Taken together, these results should further encourage researchers to utilize this probe kit regardless of tissue type or phylogenetic scale.

## Acknowledgments

The authors would like to thank the curators of APCR, EKY, KANU, LSU, MO, MT, NY, OSH, UARK, UCAC, UNA, and WAT for herbarium loans and permission to sample material for molecular analysis. Thanks also to Theo Whitsell for helpful discussions, to Jenny Hackett at the University of Kansas Genome Sequencing Core for help with sequencing, and to the staff of the Beocat computing cluster at Kansas State University for considerable help with data analysis. This study was funded by NSF award DEB 1556323 to JBB. This research was not preregistered with an analysis plan in an independent, institutional registry.

## Author contributions

JBB helped conceive the project, constructed hyb-seq libraries, performed phylogenomic analyses, and helped write the manuscript. MLM and MGZ extracted DNA and constructed genomic libraries. JRT helped conceive the project and provided taxonomic guidance. HJH constructed hyb-seq libraries. LDW performed genomic SNP analyses and helped write the manuscript. MGJ assisted with phylogenomic and genomic SNP analyses.

## APPENDIX 1

Samples included in molecular analysis. Taxa appear in bold followed by the Illumina library number (which serves as a unique identifier), collector, collector number, year collected, herbarium, country, state/province, county/parish, and GenBank accession number.

***Solidago auriculata*** – IL86, Thomas, 107076, 1988, (MO), United States, Louisiana, Caldwell,,; IL87, Thomas, 121475, 1990, (WAT), United States, Louisiana, Caldwell,,; ***Solidago brachyphylla*** – IL76, Kral, 94641, 2003, (MO), United States, Alabama, Lee,,; IL77, Anderson, 15186, 1994, (MO), United States, Florida, Okaloosa,,; IL78, Semple, 10957, 1999, (?), United States, Florida, Gadsden,,; ***Solidago delicatula*** – IL04, Nunn, 9017, 2003, (UARK), United States, Arkansas, Hempstead,,; IL08, Taylor, 16887, 1974, (KANU), United States, Oklahoma, Pushmataha,,; IL15, Morse, 8636, 2002, (KANU), United States, Oklahoma, Murray,,; IL16, Bare, 1753, 1968, (KANU), United States, Oklahoma, Lincoln,,; IL19, Thompson, S0003, 1988, (KANU), United States, Oklahoma, Comanche,,; IL20, Holland, 9736, 1999, (KANU), United States, Kansas, Neosho,,; IL21, McElderry, 281, 2005, (UARK), United States, Arkansas, Polk,,; ***Solidago drummondii*** – IL83, Hyatt, 3514.03, 1990, (MO), United States, Arkansas, Baxter,,; IL84, Hyatt, 3670.45, 1990, (MO), United States, Arkansas, Marion,,; IL85, Miller, 5493, 1990, (MO), United States, Missouri, Jefferson,,; ***Solidago fistulosa*** – IL79, Semple & Suripto, 10120, 1991, (WAT), United States, Louisiana, St. Tammany,,; IL80, Semple & Suripto, 9785, 1991, (WAT), United States, South Carolina, Clarendon,,; IL81, Semple, 10879, 1999, (WAT), United States, Georgia, Echols,,; IL82, Semple, 11624, 2006, (WAT), United States, Virginia, Southampton,,; ***Solidago patula*** – IL89, Semple & Horsburgh, 10575, 1995, (WAT), Canada, Ontario, Haldimand-Norfolk,,; IL90, Semple, 11132, 2002, (WAT), United States, North Carolina, Avery,,; ***Solidago rugosa*** – IL01, Kral, 44713B, 1971, (KANU), United States, Alabama, Butler,,; IL05, Fryxell, 3148, 1979, (MO), United States, Texas, Grimes,,;IL09, Andreasen, 46, 1970, (MO), United States, Missouri, Franklin,,; IL17, MacDonald, 9174, 1996, (MO), United States, Mississippi, Grenada,,; IL23, Nunn, 9243A, 2003, (UARK), United States, Arkansas, Yell,,; IL24, Gates, 34, 1973, (MO), United States, Louisiana, Jackson,,; IL25, Semple, 9472, 1991, (WAT), United States, New York, Livingston,,; IL26, Semple, 2399, 1976, (WAT), Canada, Ontario, Brant,,; IL30, Miller, 465, 1970, (UARK), United States, Arkansas, Lincoln,,; IL31, Holland, 10700, 2003, (KANU), United States, Arkansas, Garland,,; IL32, Semple, 10086, 1991, (WAT), United States, Louisiana, Calcasieu,,; IL58, Brant, 6323, 2007, (MO), United States, Missouri, Wayne,,; IL72, Semple & Suripto, 10122, 1991, (WAT), United States, Mississippi, Harrison,,; IL73, Semple & Suripto, 10093, 1991, (WAT), United States, Louisiana, Rapides,,; IL74, Semple & Suripto, 10086, 1991, (WAT), United States, Louisiana, Calcasieu,,; IL75, Semple & Suripto, 9564, 1991, (WAT), United States, Massachusetts, Barnstable,,; ***Solidago uliginosa*** – IL91, Semple & Keir, 4602, 1980, (WAT), Canada, Quebec, Wolfe,,; IL92, Semple & Suripto, 9576, 1991, (WAT), United States, Massachusetts, Essex,,; IL93, Semple, 9067, 1998, (WAT), United States, Minnesota, Aitkin,,; ***Solidago ulmifolia* aff. var. *palmeri*** – IL06, Davis, 3911, 1911, (MO), United States, Missouri, Marion,,; IL29, Thompson, 888, 1974, (UARK), United States, Arkansas, Newton,,; IL51, Demaree, 37672, 1955, (LSU), United States, Arkansas, Garland,,; IL52, Thomas, 125820, 1991, (OSH), United States, Arkansas, Independence,,; IL56, Morse, 3832, 1999, (KANU), United States, Arkansas, Scott,,;IL59, Leidolf, 596, 1994, (MO), United States, Mississippi, Oktibbeha,,; IL61, Taylor, 27275, 1978, (MO), United States, Missouri, Wright,,; IL64, Henderson, 95786, 1995, (MO), United States, Missouri, Benton,,; IL66, Kral, 33035, 1968, (MO), United States, Alabama, Lamar,,; IL67, McElderry, 2666, 2005, (UARK), United States, Arkansas, Monroe,,; ***Solidago ulmifolia* var.*palmeri*** – IL07, Demaree, 38132, 1955, (KANU), United States, Arkansas, Garland,,; IL10, Stephens, 17667, 1967, (KANU), United States, Nebraska, Cass,,; IL47, Bodine, 10, 1994, (MO), United States, Missouri, Wayne,,; IL48, Marisco, 3544, 2002, (UARK), United States, Arkansas, Montgomery,,; IL49, Roberts, 441, 1977, (UARK), United States, Arkansas, Hot Spring,,; IL50, Demaree, 34297, 1953, (NY), United States, Arkansas, Montgomery,,; IL53, Ray, 5843, 1955, (NY), United States, Mississippi, Clay,,; IL54, Mohr, SN, 1892, (UNA), United States, Alabama, Dallas,,; IL55, Moore, 65-267, 1965, (APCR), United States, Arkansas, Yell,, *; **Solidago ulmifolia*** **var. *ulmifolia*** – IL03, Timme, 22450, 2011, (MO), United States, Arkansas, Benton,,; IL27, McElderry, 308, 2005, (UARK), United States, Arkansas, Montgomery,,; IL28, McDaniel, 3405, 1962, (NY), United States, Mississippi, Webster,,; IL33, Semple, 9957, 1991, (MO), United States, Arkansas, Washington,,; IL34, Smith, 3986, 2004, (MO), United States, Missouri, Barry,,; IL57, Semple, 9092, 1988, (WAT), United States, Wisconsin, Winnebago,,; IL60, Ovrebo, W1056, 1989, (MO), United States, Missouri, Pulaski,,; IL62, Semple, 9893, 1991, (WAT), United States, Missouri, Wayne,,; IL63, Morse, 10562, 2004, (KANU), United States, Kansas, Linn,,; IL65, Redfearn, 32727, 1981, (MO), United States, Missouri, Christian,,; IL69, Lovejoy, s.n., 1998, (OSH), United States, Massachusetts, Hampden,,; IL70, Nunn, 7178, 2002, (UARK), United States, Arkansas, Prairie,,; IL71, Beck, 59, 1997, (EKY), United States, Kentucky, Fleming,.

